# Regulation of replicative histone RNA metabolism by the histone chaperone ASF1

**DOI:** 10.1101/2022.11.30.518476

**Authors:** Shweta Mendiratta, Dominique Ray-Gallet, Alberto Gatto, Sébastien Lemaire, Maciej A. Kerlin, Antoine Coulon, Geneviève Almouzni

## Abstract

In S phase, duplication and assembly of the whole genome into chromatin requires upregulation of replicative histone gene expression. Here, we explored a potential role of histone chaperones in this process thereby linking chromatin assembly with histone production in human cells. Depletion of the ASF1 chaperone specifically decreased the pool of replicative histones both at the levels of soluble protein and total RNA, while depletion of CAF-1 did not. Most replicative histone genes decreased in their overall expression as revealed by total RNA-seq. In contrast, both their newly synthesized RNAs and nascent RNAs at transcription sites increased as shown by 4sU-labeled RNA-seq and single-molecule RNA FISH, respectively. Further inspection of the sequences corresponding to replicative histone RNAs showed a 3’ processing defect, leading to unprocessed transcripts usually targeted for degradation. We discuss how this regulation of replicative histone RNA metabolism by ASF1 fine-tunes the histone dosage to avoid unbalanced situations deleterious for cell survival.

## INTRODUCTION

Histones are the protein components of the basic unit of chromatin, the core particle of the nucleosome, which consists of a (H3-H4)_2_ tetramer flanked by two H2A-H2B dimers around which 147 bp of DNA wraps (Luger, 1997; Zhou et al., 2019). They play a central role in defining chromatin states associated with distinct cell fates. They are classified into replicative and non-replicative/replacement histone variants. While the latter do not exhibit S phase regulation in their expression, the replicative histone variants show a major peak in expression early during S phase to support chromatin assembly during replication of the genome (Armstrong and Spencer, 2021; Marzluff et al., 2008; Mendiratta et al., 2019). In addition, beside distinct regulation in expression timing, replicative and non-replicative histone genes also differ in their genomic organization and RNA structure. In most metazoans, the replicative histone genes are organized in clusters, lack introns (Marzluff et al., 2008; Marzluff and Koreski, 2017; Mendiratta et al., 2019) and are expressed in specialized nuclear bodies known as histone locus bodies (HLBs) (Hur et al., 2020; Nizami et al., 2010). Their mRNAs do not contain a polyA tail but present a 3’conserved stem loop (SL) structure (Marzluff et al., 2008; Marzluff and Koreski, 2017). These distinct features enable to provide the necessary increase in histone supply for nucleosome assembly during genome duplication in S phase with a ∼40-fold increment in histone mRNA levels during the G1/S phase transition. This involves the upregulation of both transcription and 3’ end processing (Harris et al., 1991). The expression of replicative histone genes is therefore tightly regulated during the cell-cycle both transcriptionally and post-transcriptionally and involves a number of actors (Marzluff et al., 2008; Marzluff and Koreski, 2017; Mendiratta et al., 2019).

From the time of their synthesis in the cytoplasm to their delivery into the chromatin or after eviction, histones are always escorted by histone chaperones with dedicated functions (Gurard-Levin et al., 2014; Hammond et al., 2017). This prevents free histones, that are positively charged, to engage into promiscuous interactions with any acidic partner, notably with DNA, that could form aberrant aggregates in the cell. During replication in human cells, two main chaperones ensure the deposition of H3-H4 onto DNA: Chromatin assembly factor 1 (CAF-1) which comprises three subunits (p150/CHAF1A, p60/CHAF1B and p48) (Moggs et al., 2000; Smith and Stillman, 1989) and Anti-silencing factor 1 (ASF1), with two paralogs in mammals ASF1a and ASF1b (Abascal et al., 2013) and originally identified in yeast (Le et al., 1997). Interestingly, on the one hand, ASF1 binds the newly synthesized replicative histones H3.1/H3.2-H4 (Tagami et al., 2004) to hand them off to the downstream chaperone, CAF-1, for deposition onto the duplicated DNA strands in a DNA synthesis-coupled (DSC) manner (Mello et al., 2002; Tyler et al., 1999). On the other hand, ASF1 also promotes the recycling of parental histones during replication (Clément et al., 2018; Groth et al., 2007). In addition, ASF1 binds the non-replicative variant H3.3 and hands it off to the downstream chaperone Histone regulator A (HIRA) for deposition of H3.3 in a DNA synthesis independent (DSI) manner (Ahmad and Henikoff, 2002; Tagami et al., 2004; Tang et al., 2006). Finally, in human cells, ASF1 but not CAF-1, also provides a buffering system for histone excess generated in response to stalled replication (Groth et al., 2005) indicating yet another role for ASF1 in regulating the flow of replicative histones in higher eukaryotes. However, to date, roles of these chaperones in histone RNA metabolism in mammals had remained unexplored. This is particularly interesting to consider given that in budding yeast, where there are no distinct replicative and non-replicative H3 variants, the single ASF1 ortholog which synergizes with CAF-1 in histone deposition during replication (Sharp et al., 2001; Tyler et al., 1999) participates in activating transcription of histone genes in S phase and transcriptional repression outside S phase in combination with Hir1, the budding yeast counterpart of HIRA (Sutton et al., 2001). We thus decided to explore how the key histone chaperones involved in DNA synthesis-coupled chromatin assembly could contribute to the critical regulation of expression of replicative histone genes in human cells during S phase. We first explored in human cells the effect of depleting either ASF1 or CAF-1 on the protein and RNA levels of replicative histones. Strikingly, ASF1 depletion, but not CAF-1 (p60 subunit) depletion, leads to a specific decrease of replicative histones at the levels of soluble protein and total transcript, while those corresponding to non-replicative histones remained unaffected. From total RNA extracted from asynchronous and synchronized human cells, we performed RNA-seq and found that most of the annotated replicative histone genes decreased in expression upon ASF1 depletion during S phase. However, by single-molecule RNA FISH we found an increase in replicative histone nascent transcripts at transcription sites and by 4sU- labeled RNA-seq we also detected an increase in newly synthesized replicative histone transcripts. These findings indicate that the decrease in expression of replicative histone genes in ASF1-depleted cells cannot be due to a decrease at the level of transcription. We then inspected closely the sequences at the 3’ end of the replicative histone transcripts in our RNA-seq data and detected a defect of their 3’ processing. Thus, we propose that in mammals ASF1 plays a role in the unique regulation of replicative histone RNA metabolism.

## RESULTS

### ASF1 depletion, in contrast to CAF-1 depletion, reduces replicative histones at the levels of both soluble protein and total RNA

To explore roles of ASF1 or CAF-1 in the regulation of the expression of replicative histone genes, we analyzed the effect of their respective depletion on replicative histones both at protein and RNA levels. We transfected HeLa cells with siRNAs targeting ASF1a and ASF1b (siASF1), the p60 subunit of CAF-1 (sip60) or control (siCtrl). We depleted both ASF1a and ASF1b paralogs, since they most often compensate the depletion of the other (Abascal et al., 2013; Gao et al., 2018). 48 h post-transfection, we harvested cells and isolated total RNAs or proteins in cytosolic, nuclear and chromatin fractions (Figure 1a). In the chromatin fraction, the replicative histones H3.1 and H3.2 (recognized simultaneously by the same antibody) decreased upon both ASF1 and p60 depletions as shown by Western blot analysis (Figure 1b). This decrease is consistent with their role in promoting the deposition of these histones coupled to DNA synthesis during replication. Strikingly, in the nuclear and especially cytosolic soluble extracts, the replicative histones H3.1/3.2 and H4 significantly decreased upon ASF1 depletion (a 3-fold to 4- fold decrease in the cytosolic extract) (Figure 1c and 1d). In contrast, they rather increased upon p60 depletion likely reflecting deficiency in their deposition. This behaviour of replicative histones differs from non-replicative H3.3 which amounts of soluble protein did not change upon both ASF1 and p60 depletions (Figure 1c and 1d). We confirmed these findings by using different siRNAs targeting ASF1a and b or p60 (Supplementary figure 1a and 1b). Furthermore, we also noted that soluble replicative histone H2B also decreased (Supplementary figure 1c), thus ASF1 depletion may in fact affect all types of replicative histones in a coordinated fashion.

**Figure 1.**
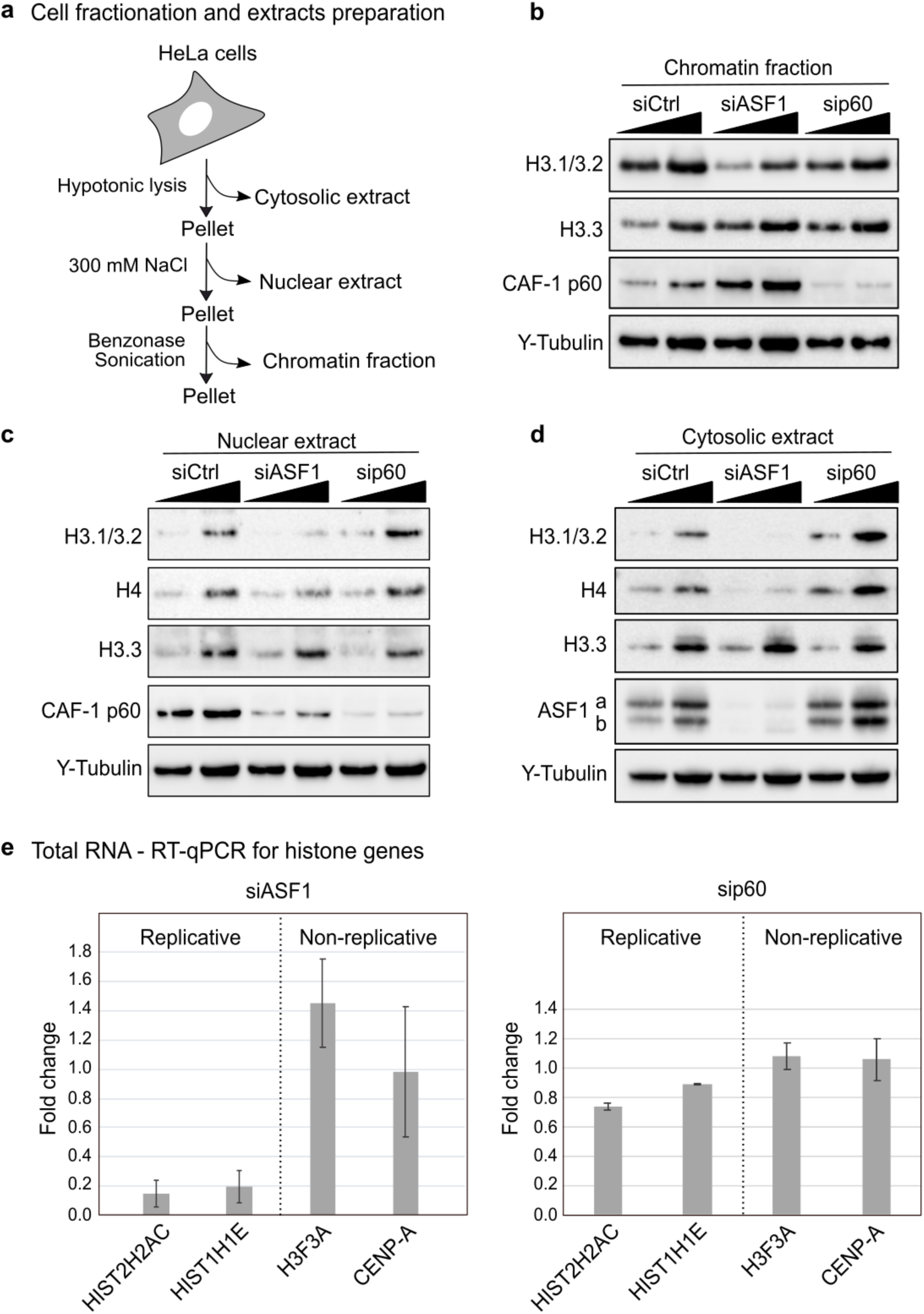
ASF1 depletion reduces replicative histones at the levels of both soluble protein and total RNA. **(a)** Scheme depicting the preparation of cytosolic, nuclear and chromatin fractions from HeLa cells. **(b, c, d)** Western blot analysis of chromatin fraction (b), nuclear extract (c) and cytosolic extract (d) from HeLa cells harvested 48 h after siRNA transfection (siCtrl, siASF1 or siCAF-1-p60). Two amounts 1x and 2x were loaded. The replicative histones H3.1/3.2 and H4 and the non-replicative histone H3.3 were examined, γ-tubulin was used as loading control. ASF1a and b were shown in cytosolic extracts in which they are the more abundant. CAF-1 p60 was not shown in the cytosolic extracts as it is not detected in this fraction. The distribution of CAF-1 p60 changed upon ASF1 depletion with an increase in chromatin fraction and a decrease in nuclear extract) **(e)** RT-qPCR analysis of histone RNA levels. Total RNA was isolated from HeLa cells harvested 48 h after siRNA transfection (siCtrl, siASF1 or siCAF-1-p60). Individual histone genes are the replicative HIST2H2AC (H2A type 2-C) and HIST1H1E (H1.4) and the non-replicative H3F3A (H3.3) and CENP-A. The fold change in transcript levels compared to siCtrl corresponds to the mean of three independent biological replicate experiments.

The reduction in the pool of soluble replicative histones upon ASF1 depletion prompted us to examine whether this change relates to effects at an RNA level. We thus performed RT-qPCR using total RNAs to evaluate changes in the amount of replicative histone transcripts upon ASF1 and p60 depletions. Total RNA for the replicative histone genes, HIST2H2AC and HIST1H1E strongly decreased in ASF1-depleted cells which is not the case in p60-depleted cells (Figure 1e). In contrast, total RNA did not decrease for known non-replicative histones like H3F3A and CENP-A (Figure 1e). The different behaviour after depletion of either ASF1 or CAF-1 is unlikely a consequence of the S phase progression defect, since both of them similarly affect S phase progression (Groth et al., 2007, 2005; Hoek and Stillman, 2003). We further confirmed our findings in another human cell line, U2OS (Supplementary figure 1d). Moreover, by expressing Strep-tagged ASF1a and b in siASF1-depleted HeLa cells, we could rescue the defect in replicative histones at both RNA and soluble protein levels (Supplementary figure 1e). The rescue argues for a direct importance of ASF1 and not a partner that could have been affected by the depletion.

Our first results support a specific role for ASF1, and not for CAF-1, in modulating the levels of both total RNA and soluble protein corresponding to replicative histone genes.

### ASF1 depletion reduces the expression of most annotated replicative histone genes at a transcriptome level during S phase

To analyze more broadly how ASF1 depletion affects the transcriptome and in particular the transcription of replicative histone genes, we performed RNA-seq to compare total RNA from siCtrl and siASF1-transfected cells in asynchronous cells (Figure 2a). Differential gene expression analysis revealed global transcriptional changes upon ASF1 depletion (Figure 2a and Supplementary figure 2a and 2b). Pathways associated with cell cycle progression, most notably MYC-driven pathways, ranked among the top down-regulated. Instead, among the top up- regulated pathways, we found tumor necrosis factor alpha (TNFA) signaling and the p53 pathway involved in cell cycle arrest. Interestingly, the non-replicative histone gene H1F0 was also upregulated, and it is normally reduced in cells with long-term self-renewal ability and tumorigenic potential (Morales Torres et al., 2020; Torres et al., 2016). Altogether, these observed changes are in line with the reduced proliferation and slow S phase completion observed in ASF1-depleted cells (Groth et al., 2007, 2005). A general trend that caught our attention was the noticeable reduction in the expression level of replicative histone genes upon ASF1 depletion, with most replicative histones ranking among the top down-regulated genes (Figure 2a). To be able to compare G1/S arrested and S phase cells, we then prepared total RNA from synchronized cells to generate RNA-seq data (scheme Figure 2b). Hierarchical clustering of differentially expressed genes at different S phase stages, identified six main groups of genes (annotated as 1 to 6) upon ASF1 depletion (Figure 2b, 2d and Supplementary figure 2d)). Down- regulated genes fell in groups 1, 2 and 3, while up-regulated ones in groups 4, 5 and 6. Groups 1 and 2 comprised the top down-regulated genes after ASF1 depletion and showed the strongest changes with S phase progression under control conditions (Figure 2d). Group 1 showed enrichment in genes involved in DNA replication (Supplementary table 2). Most interestingly, Group 2 stood out as it exclusively comprised replicative histone genes with the largest differences in expression along S phase progression and after ASF1 depletion (Figure 2c and 2d). Indeed, under control conditions, replicative histone genes displayed their lowest expression levels in G1/S cells and strongest induction after release into S phase, peaking gradually from early (2 h) to mid S phase (5 h). ASF1 depletion led to a major repression of all these genes, with lowest baseline levels in G1/S and a striking failure to induce their expression after release into S phase. On average, we estimated that replicative histone genes decreased by ∼2-fold, ∼3-fold and ∼6-fold relative to control at 0 h, 2 h and 5 h, respectively. Additionally, we compared the expression profile of all replicative histone genes with each other (Supplementary figure 2e) and with non-replicative histone genes (Supplementary figure 2f). Interestingly, for the few replicative histone genes not affected by ASF1 depletion they did not show a strong expression upon S phase entry in control cells, indicating that they are under separate regulatory means even if they are localized in the histone gene clusters. Thus, the regulation revealed here may not use a global local subnuclear means which would equally affect the whole locus.

**Figure 2.**
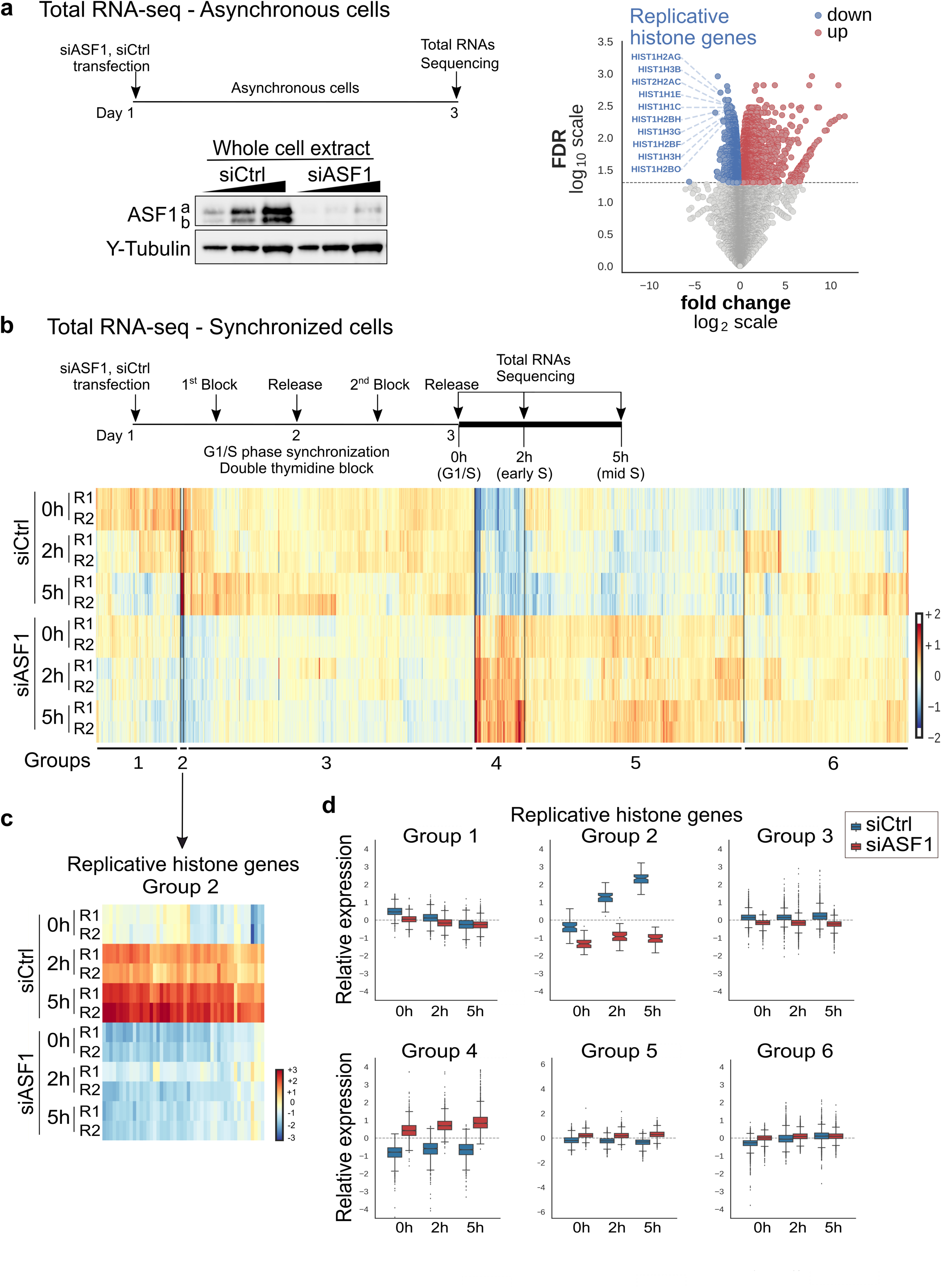
ASF1 depletion reduces the expression of most of the replicative histone genes at the level of total RNA during S phase (a) (Left top) Scheme showing the steps performed for the RNA-seq analysis from asynchronous cells. Hela cells were transfected with siRNA (siCtrl or siASF1) and after 48 h total RNA was purified and sequenced. **(Left bottom)** Western blot analysis of total extract from siCtrl- and siASF1-treated cells showing the efficiency of ASF1 depletion. **(Right)** Volcano plot highlighting the decrease of replicative histone gene expression upon ASF1 depletion from asynchronous cells. For all genes expressed in at least two samples, the plot shows the statistical significance (y-axis, - log_10_ FDR) and difference in total RNA (x-axis, log_2_ fold change) between siASF1 and siCtrl conditions. Genes showing significant differences at 5% FDR are shown in blue, for down- regulated, and red, for up-regulated. The top 10 replicative histone genes showing significant differences are highlighted in blue, ranked by adjusted p-value (- log_10_ FDR). **(b) (Top)** Scheme showing the steps performed for the RNA-seq analysis of synchronized cells. Hela cells were transfected with siRNA (siCtrl or siASF1), then synchronized using a double thymidine block protocol. The cells were harvested at 0 h (G1/S) and at 2 h and 5 h (S phase) following their release from the second thymidine block. Total RNA was purified and sequenced. **(Bottom)** Heat map showing hierarchical clustering of samples (rows) and differentially expressed genes (columns) between siCtrl and siASF at different time points after synchronization and release into S phase, including all genes showing significant differences at FDR < 0.05. The color gradient is proportional to the expression of each gene in a given sample, relative to their average across all samples (log_2_ normalized counts, mean-centered): from blue (lower than average) to red (higher than average). Six groups of genes, identified based on hierarchical clustering results (top six column clusters), are highlighted at the bottom of the heat map. The experiment was performed in duplicate, replicate 1 (R1) and replicate 2 (R2). **(c)** Heatmap showing hierarchical clustering of Group 2 which consists only of replicative histone genes, in siCtrl and siASF1 conditions. Color gradient shows expression relative to average as in 2b (log_2_ normalized counts, mean-centered): from blue (lower than average) to red (higher than average). **(d)** The six groups of genes identified by hierarchical clustering from differential gene expression between siASF1 and siCtrl in synchronized HeLa cells. For each group of genes, the box plots show the distribution of relative expression levels across all genes in siCtrl (in blue) and siASF1 (in red) conditions at 0 h (G1/S), 2 h (early S) and 5 h after release from the G1/S block (log_2_ normalized counts, mean-centered across all samples and averaged by experimental condition). Group 2 contains only replicative histone genes. See Supplementary figure 2d for Gene ontology (GO) annotations for the other groups.

In conclusion, ASF1 depletion led to global coordinated changes in transcription with a strong effect on S phase-dependent genes, among which remarkably most of the annotated replicative histone genes.

### ASF1 depletion increases replicative histones at the levels of both newly synthesized RNA and nascent RNA at transcription sites

Since the expression of replicative histone genes in mammalian cells is controlled at multiple levels (Marzluff and Koreski, 2017; Mendiratta et al., 2019), we examined first whether ASF1 plays a role at the level of their transcription. To analyse the newly transcribed RNAs, we performed 4-Thiouridine-labeled RNA-sequencing (4sU-labeled RNA-seq). To our surprise, we observed more newly synthesized RNAs for the replicative histone genes upon ASF1 depletion (Figure 3a and Supplementary figure 3a). Indeed, this was unexpected based on the decrease in their expression observed by RNA-seq and RT-qPCR. Thus, we used another approach to analyse the transcription of replicative histone genes. We employed single-molecule RNA FISH imaging to investigate the levels of nascent transcripts of replicative histone genes at transcription sites upon ASF1 depletion. Achieving single-molecule sensitivity in FISH requires targeting of multiple fluorescent probes against a transcript of interest. Replicative histone genes are relatively short and lack introns, which together makes it difficult to visualize nascent transcription events of a single histone gene. Therefore, we labeled transcripts of four replicative histone genes – *HIST1H4C*, *HIST1H1E*, *HIST1H2BF* and *HIST1H4E* – clustered within ∼100 kb on chromosome 6 (corresponding to cluster 1) (Figure 3b). When imaging transcription of the histone gene cluster, we observed RNA spots of monodispersed intensity corresponding to single RNA molecules, mostly localized in the cytoplasm, as well as a few bright spots localized exclusively in the nucleus, corresponding to the transcription sites of the histone gene cluster (Figure 3b). Our observation of a maximum of 3 transcription sites per nucleus corresponding to cluster 1 (Supplementary figure 3b) is in line with the number of histone locus bodies identified in Hela cells by DNA-FISH (Ghule et al., 2009). Upon ASF1 depletion, the intensity of transcription sites of replicative histones increased by 78% compared to the control condition (Figure 3c and supplementary figures 3c and 3d) while the transcription of the *FOS* gene, used as a control, did not change (Supplementary figure 3e). In contrast, we did not detect any change in intensity at transcription sites of replicative histones upon CAF-1 (p60) depletion (Figure 3c).

**Figure 3.**
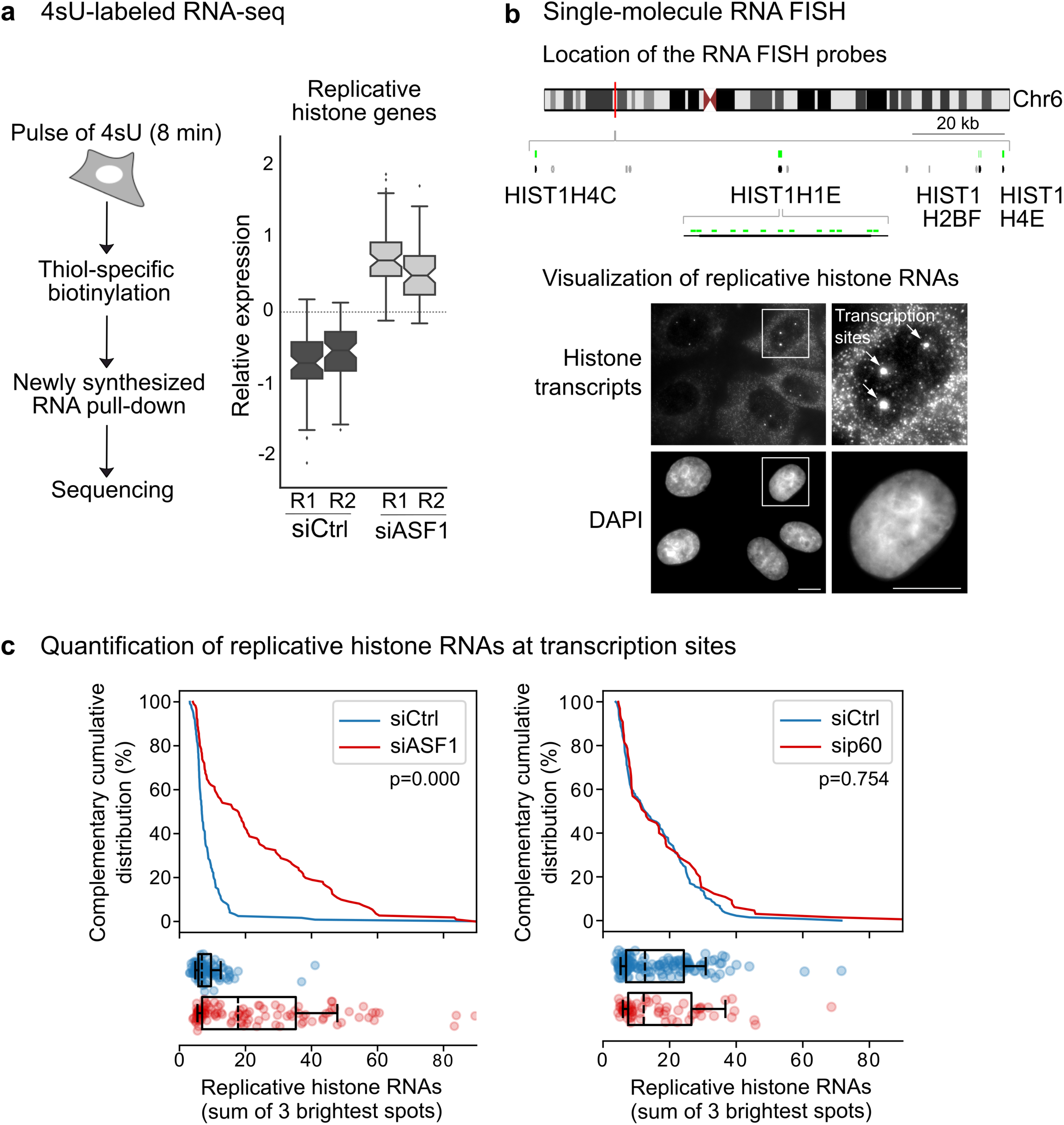
ASF1 depletion increases replicative histones at the levels of both newly synthesized RNA and RNA at transcription sites (a) (Left) Scheme showing the procedure for 4sU-labeled RNA sequencing (Fuchs et al., 2015) **(Right)** Relative expression of replicative histone genes from transcripts in control condition (siCtrl) and upon ASF1 depletion (siASF1). For each 4sU-labeled RNA-seq replicate, the box plot shows the distribution of relative expression levels for all replicative histone genes in siCtrl and siASF1 conditions (mean-centered counts, TMM-normalized and log_2_ transformed). **(b) (Top)** Scheme of chromosome 6 illustrating the position of single-molecule RNA FISH probes (green). The four labeled replicative histone genes are depicted in black. Grey dots correspond to other histone genes located in the locus. **(Bottom)** Representative image of labeled replicative histone transcripts. Single histone RNAs are visible as low-intensity fluorescence spots. Sites of transcription, indicated by white arrows, are visible as bright spots and found exclusively inside nuclei. Nuclei stained with DAPI are shown in blue. Scale bar 10 μm **(c)** Complementary cumulative distributions of the number of RNAs at transcription sites per nucleus (calculated as the sum of the 3 brightest spots) in siCtrl and siASF1 conditions (Left) and in siCtrl and sip60 CAF-1 conditions (Right). Each plot represents a single biological replicate. See Supplementary figure 3d for other replicates for siCtrl and siASF1 conditions. The p-values of Kolmogorov-Smirnov test are indicated. Scatterplots show the numbers of replicative histone RNAs at transcription sites per nucleus in control condition (siCtrl, blue) and upon ASF1 or p60 depletions (siASF1 or sip60, red). Dashed lines show the median RNA numbers, boxes show the top and bottom quartiles and whiskers show the 10- and 90-percentile.

In conclusion, while ASF1 depletion led to a decrease in total transcripts corresponding to replicative histones, our latter results showed that when considering newly synthesized RNAs and nascent RNAs at sites of transcription, their amounts are not reduced and even increased. While this was at first counterintuitive, given the multiple levels involved in regulating replicative histone gene expression in mammals, to understand how replicative histone RNAs change one must also inspect the nature of nascent transcripts, and their processing. We thus explored whether ASF1 may impact the 3’ processing of replicative histone pre-mRNAs.

### ASF1 depletion affects the 3’ processing of replicative histone pre-mRNAs

The peculiar feature of replicative histone transcripts in mammals, that are not polyadenylated, is associated with a unique post-transcriptional regulation involving a series of factors (Supplementary figure 4a) (Marzluff et al., 2008). To ask whether ASF1 depletion could affect the 3’ processing of replicative histone transcripts, we first wonder whether depleting known factors involved in this 3’ processing may mimic the effect of ASF1 depletion on replicative histones. We depleted therefore either FLice-associated huge protein (FLASH) (Yang et al., 2009) or Stem-loop binding protein (SLBP) (Wang et al., 1996) by siRNA and analyzed the impact on soluble replicative histones. Strikingly, by Western blot analysis of cytosolic extracts, we observed a comparable decrease in replicative H3.1/3.2 and H4 without affecting the non-replicative H3.3 in siFLASH-, siSLBP- and siASF1-transfected cells (Figure 4a). The apparent decrease of ASF1a in FLASH- and SLBP-depleted cells is likely due to the increase of slower migrating phosphorylated ASF1a forms as previously observed in cells experiencing deficiency of new replicative histones notably by FLASH depletion (Klimovskaia et al., 2014). Moreover, remarkably, in the chromatin fraction, the amounts of replicative histone H3.1/3.2 decreased upon FLASH and SLBP depletions in a manner that compares to our observations upon ASF1 depletion (Supplementary figure 4d). This is typically expected for factors required for regulating the expression of replicative histone genes. Both ASF1 and FLASH depletions resulted in an increase of SLBP in cytosolic, nuclear and chromatin fractions (Figure 4a and Supplementary figure 4b). This increase in SLBP, not observed in CAF-1 (p60)-depleted cells (Supplementary figure 4c), argues that the increase of SLBP upon ASF1 depletion is not merely due to the accumulation of cells in S phase. We also examined the Sm-like protein LSM10, another specific actor of the replicative histone pre-mRNAs 3’ processing which is part of the U7snRNP complex and did not detect significant change for this protein (Supplementary figure 4b). The comparable effects observed in siASF1-, siFLASH and siSLBP-transfected cells suggest that ASF1 depletion may impact the post-transcriptional regulation of replicative histone genes in a similar manner.

**Figure 4.**
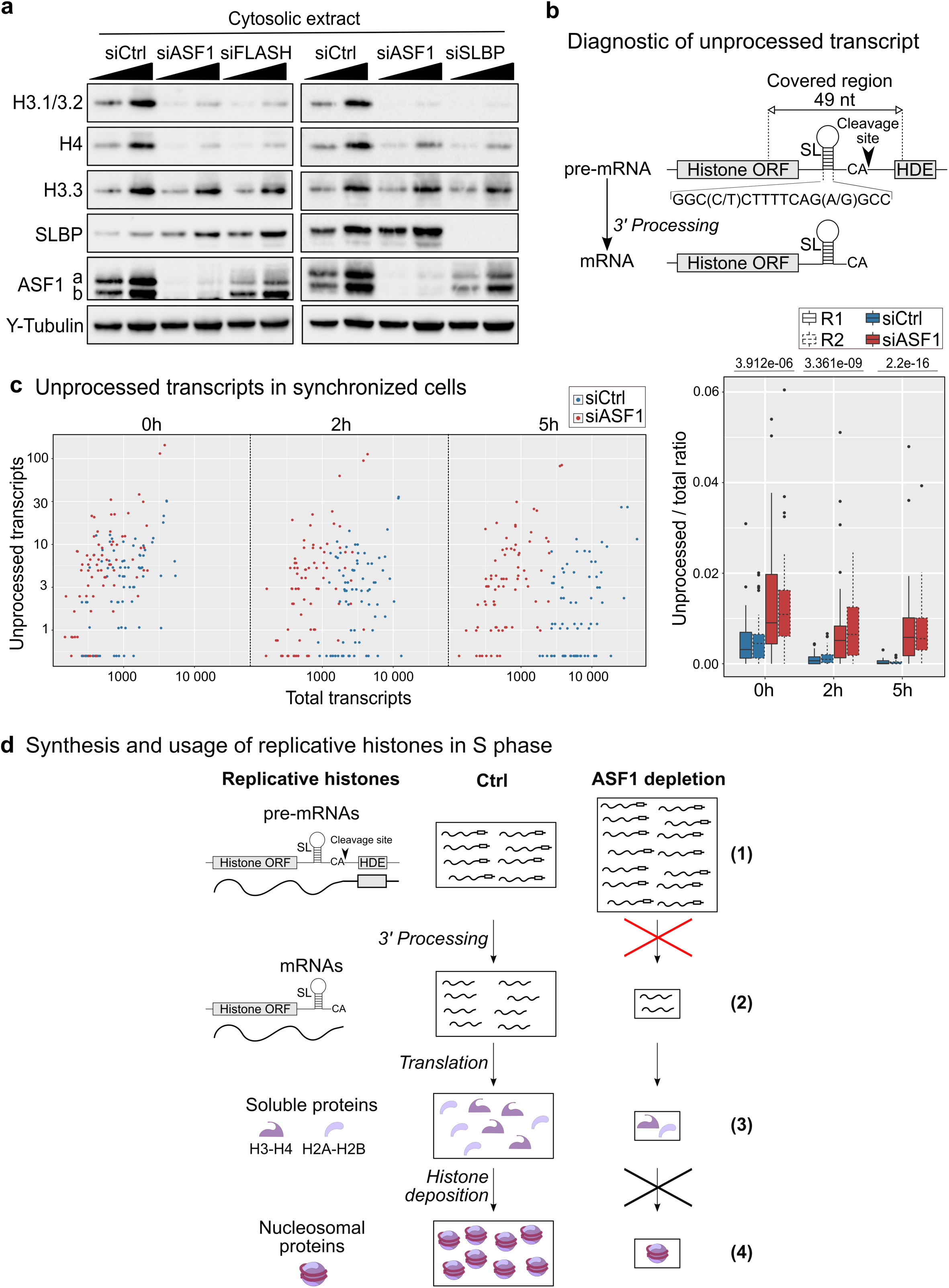
ASF1 depletion affects the 3’ processing of replicative histone transcripts. **(a)** Western blot analysis of cytosolic extract from HeLa cells harvested 48 h after siRNA tranfection (siCtrl, siASF1, siFLASH or siSLBP). Two amounts 1x and 2x were loaded, γ-tubulin was used as loading control. **(b)** Scheme showing the structure of replicative histone pre-mRNA and mRNA after 3’ processing. The cleavage occurs downstream the Stem-Loop (SL) and upstream the Histone Downstream Element (HDE). The indicated region of 49 nucleotides in an RNA-seq fragment is diagnostic of an unprocessed replicative histone transcript. This fragment encompasses the cleavage site, starting 20 nucleotides upstream of the SL (16 nt long) and ending 13 nucleotides downstream of the SL, reaching the HDE. **(c)** Comparison of the amounts of unprocessed transcripts between siASF1 and siCtrl conditions using our RNA-seq data of synchronized cells (at times 0 h, 2 h and 5 h after release). **(Left)** The replicative histone genes are plotted according to their normalized number of unprocessed and total transcripts. Each dot is one replicative histone gene. Genes with a null number of unprocessed transcripts are displayed at the value of 0.5. **(Right)** The box plots show the ratio of the number of unprocessed transcripts to total transcripts (total number of reads mapping in each annotated replicative histone gene). The results for the two replicates R1 and R2 are shown. We found ∼2.5, ∼5.7 and ∼20.4 times more unprocessed transcripts in siASF1 as compared to siCtrl at 0 h, 2 h and 5 h post-release, respectively. The p-values are indicated. **(d)** Model showing the impact of ASF1 depletion on the synthesis and usage of replicative histones during S phase. ASF1 depletion leads to an increased transcription of replicative histone genes and the accumulation of their newly synthesized pre-mRNAs that cannot be processed and likely targeted to degradation (1). The lack of mature mRNAs due to a defect of 3’ processing (2) leads to reduced levels of soluble replicative histones (3) and ultimately a decrease of nucleosomal replicative histones (4) due to the decrease of both synthesis and usage (histone deposition) of replicative histones.

The 3’ processing of replicative histone pre-mRNAs, with a cleavage occurring between the stem loop (SL) and the Histone Downstream Element (HDE), produces mRNAs containing the SL and deprived of the sequence downstream the cleavage site and notably the HDE (Figure 4b) (Marzluff et al., 2008). We examined a role of ASF1 in the 3’ processing of replicative histone pre-mRNAs, taking advantage of our 4sU-labeled RNA-seq and total RNA-seq data. To define unprocessed transcripts, we selected them based on a sequence of about 49 nucleotides that encompasses the cleavage site, starting 20 nucleotides upstream of the SL and ending 13 nucleotides downstream of the SL to reach the HDE (Figure 4b). Using our sequencing data, we estimated the relative amounts of unprocessed transcripts by computing the number of fragments mapping to the non-cleaved 3’ region of each replicative histone gene (as defined above) to the total number of fragments mapped to the same gene. From the 4sU-labeled RNA-seq, we did not detect more unprocessed transcripts upon ASF1 depletion, but we found more total transcripts (Supplementary figure 4f) suggesting that the increase of newly synthesized RNAs for replicative histone genes in ASF1-depleted cells (Figure 3a) reflects mainly an increase of transcription. However, from the total RNA-seq in synchronized cells, we could reveal ∼ 1.9, 4.9 and 14 times more unprocessed transcripts upon ASF1 depletion as compared to control condition at 0 h, 2 h and 5h post-release, respectively (Figure 4c). This increasing difference reflects that, while the fraction of unprocessed transcripts decreased strongly over time in the control cells, this fraction only decreased moderately in ASF1-depleted cells. From the total RNA-seq in asynchronous cells, we found a same trend with ∼ 2.7 times more of unprocessed transcripts in ASF1-depleted cells as compared to control cells (Supplementary figure 4e).

We conclude that ASF1 depletion, while increasing the transcription of replicative histone genes, affects the 3’ processing of their pre-mRNAs.

## DISCUSSION

Here, we explored whether the key H3-H4 histone chaperones, ASF1 and CAF-1, both involved in DNA synthesis-coupled chromatin assembly, could specifically control the provision of replicative histone RNA and protein with their usage in S phase. We discovered a unique role for ASF1 in regulating specifically the expression of most annotated replicative histone genes as illustrated in figure 4d (see also Supplementary figure 4g for CAF-1 depletion). We revealed that ASF1 depletion led to reduced amounts of replicative histones both at total RNA and protein levels while the amounts of both newly synthesized RNAs and nascent RNAs detected at transcription sites increased. Importantly, although this result was at first counterintuitive, we found that ASF1 depletion affects the 3’ processing of replicative histone pre-mRNAs resulting in a decrease of mature mRNAs and therefore consequently of replicative histone proteins. Our data thus argue for a new role of mammalian ASF1 in the control of the replicative histone gene expression by regulating their RNA metabolism.

To explain the apparent opposite effects on replicative histone RNA metabolism that we observed upon ASF1 depletion, with more newly synthesized RNAs but less total RNAs, we propose two scenarios that are not mutually exclusive. In the first one, ASF1 depletion activates the transcription of replicative histone gene expression. This is supported by the fact that the ASF1a and ASF1b counterpart in yeast contributes to the transcriptional repression of three of the four histone gene pairs outside of S phase (Sutton et al., 2001). This deregulated transcription would then in turn induce a defect of the 3’ processing of replicative histone pre-mRNAs. Indeed, 3’ processing defects occur when transcription is perturbed (Narita et al., 2007; Saldi et al., 2018). In the second scenario, ASF1 depletion would affect first the 3’ processing of replicative histone pre-mRNAs. This possibility is supported by the identification of ASF1 in a screen for new factors involved in replicative histone pre-mRNA processing in Drosophila (Wagner et al., 2007). In this case, the increase of transcription may reflect a compensatory mechanism when mature replicative histone mRNAs become limiting. The increase of the amount of SLBP in both ASF1 and FLASH depletions could also be part of this compensatory mechanism. Moreover, the fact that depleting key processing factors specific of replicative histone pre-mRNAs like FLASH and SLBP leads to phenotypes that compares with those observed after ASF1 depletion further connects ASF1 to the replicative histone mRNA processing. Indeed, FLASH or SLBP depletions, both lead to a decrease in replicative histones, S phase progression and ssDNA formation defects in response to replication inhibitors as observed in cells lacking ASF1 (Mejlvang et al., 2014). In both scenarios, unprocessed replicative histone transcripts accumulate, while the levels of total transcripts are reduced, this suggests a targeted degradation of unprocessed pre-mRNAs as reported (Romeo et al., 2014). Future work should dissect further how and where ASF1 intervene in the RNA metabolism specific of the replicative histones.

Previous reports showed that both ASF1 and CAF-1 depletions affect S phase progression due to defects in nucleosome assembly during replication (Groth et al., 2007, 2005; Hoek and Stillman, 2003). We should now integrate how the effect on the expression of replicative histone genes in ASF1-depleted cells could also contribute to the defect of S progression. This is particularly important given that a lack in replicative histones can prevent S phase progression (Barcaroli et al., 2006; Mejlvang et al., 2014; Nelson et al., 2002). Of note, arrests in S phase progression upon overexpression of the histone chaperone HIRA had been linked to defect in the expression of replicative histone genes (Nelson et al., 2002). While at first, these observations had led to propose that HIRA acted as a repressor of replicative histone genes, as reported in yeast (Sherwood, 1993; Spector et al., 1997), alternative explanations could build on our findings. Indeed, overexpression of the HIRA mutant that does not interact with ASF1 (Tang et al., 2006), cannot interfere with the expression of replicative histone genes. Thus, considering our new findings, the difference between overexpression of HIRA wild type and HIRA lacking ASF1 interaction, could in fact reflect the trapping of ASF1 by HIRA, preventing ASF1 to play its role in stimulating the replicative histone gene expression. More generally, we believe that the unique role of ASF1 in the synthesis of replicative histones offers a different framework for the interpretation of other previous observations linked to ASF1 defects. Indeed, most transcriptomic data have been generated from poly(A+) RNAs and do not include replicative histone RNAs, and thus a number of studies probably missed the role of ASF1 on the expression of replicative histone genes. Notably, the up-regulation of ASF1a or ASF1b observed in various cancers (Abascal et al., 2013; Ray-Gallet and Almouzni, 2022) that gives tumoral cells a proliferation advantage, may actually relate to a combined stimulation of both replicative histone synthesis and deposition processes, an aspect under-explored in tumoral material so far. We postulate that fine tuning ASF1 dose and function would thus provide the cell with an ingenious way to prevent imbalance in histone provision. This would be key to deal either with excess of unused and non-chaperoned replicative histones or decrease histone provisions that would compromise establishing a proper nucleosome density, both situations that could be deleterious for the cell.

## EXPERIMENTAL PROCEDURES

### Cell lines, expression plasmids, siRNA and transfections

HeLa and U2OS cells were grown in DMEM 1X 4.5 g/L D-Glucose, L-Glutamine, Pyruvate (Gibco) with 10% Fetal Cow Serum FCS (Eurobio) and 1% Penicillin/Streptomycin (Gibco). For cell counting, cells were trypsinized and counted using Vi-Cell XR Cell Viability Analyzer (Beckman coulter). Cells were transfected with siRNA at 100 nM concentration using Lipofectamine RNAiMAX (Invitrogen, 13778-150) as previously shown in (Clément et al., 2018) and for plasmids using Lipofectamine 2000 (Invitrogen, 11668-019). Sequences used for the siRNA are mentioned in the supplementary table 1. Plasmids pEXPR-IBA105-ASF1a and - ASF1b expressing Onestrep-ASF1a and Onestrep-ASF1b, respectively were described previously (Groth et al., 2007).

### Cell staining and FACS

To analyse the cell cycle profile, around one million cells per condition were harvested using trypsin and fixed with 70% cold ethanol. Cells were then stained with propidium iodide using FxCycle^TM^ PI/RNase solution (ThermoFisher Scientific, F10797), according to the manufacturer’s instructions. The data acquition was done using FACS Attune NxT machine (ThermoFisher Scientific) and cell cycle profile was assessed with FlowJo software (V10.1r5).

### 4sU-labeled RNA-seq and total RNA-seq

For the 4sU-labeled RNA-seq, cells were cultured in 15 cm dish and 10 ml of new medium containing 1 mM 4sU was added for 8 minutes (Fuchs et al., 2015). After completion of 8 minutes, cells were lysed, and total RNA was extracted directly from the cell culture dishes using the RNeasy Mini Kit according to the manufacturer’s instructions. The total RNA was DNase treatment using the TURBO DNA-free kit (ThermoFisher Scientific, AM1907) using manufacturer’s instructions. The quantity of isolated RNA was measured using the Nanodrop and quality was checked by Tapestation. For 4sU-labeled RNA purification, 100-200 μg RNA was used for RNA biotinylation and purification was performed using Streptavidin-coupled Dynabeads (Fuchs et al., 2015). 10 ng of total and 4sU-labeled RNA was used for library preparation using SMARTer Stranded Total RNA-seq kit and sequencing was performed by the next-generation sequencing platform at Institut Curie.

### RNA-Seq analysis

Reads were aligned to the human reference genome (GRCh38 assembly) based on Ensembl gene annotation (release 95) with hisat2 (version 2.1.0), run in paired-end mode with default parameters. Gene-level counts were computed from primary alignments with MAPQ > 2 using featureCounts (Subread package version 1.6.3) in paired-end mode with the -s 2 option for reverse-stranded libraries. Raw counts were normalized for differences in library size (counts per million) and across samples (via trimmed mean of M-values normalization, TMM) using edgeR (version 3.28.0). Differential expression analyses were performed with edgeR. We assessed the effect of ASF1 depletion in asynchronous cells by fitting a quasi-likelihood negative binomial generalized log-linear model (GLM) comparing siASF1 to control. To evaluate the effect of ASF1 depletion in synchronized cells before (0 h) and after release into S phase (2 h), we similarly fitted a GLM including both additive and synergistic terms to model the stand-alone effect of S phase progression (changes in expression at 2 h and 5 h relative to 0 h) and ASF1 depletion (siASF1 relative to control across all time points), as well as at the specific effect of ASF1 depletion at different time points after release into S phase (siASF1 relative to control at 2 h and 5 h). Full details about the variable encoding and GLM formulation are provided in Supplementary table 2 (sheet 1), along with the TMM-normalized counts for both asynchronous (sheet 2) and synchronous cells (sheet 3). For each effect, we applied a quasi-likelihood F-test on the respective GLM coefficient to assess its significance, followed by multiple testing correction via the Benjamini-Hochberg method (Supplementary table 2, sheet 4 for asynchronous cells, sheet 5 for synchronous cells). A false discovery rate (FDR) cut-off of 0.05 was used to identify differentially expressed genes (DEGs). For hierarchical clustering analyses in synchronous cells, we tested the combined effect of all coefficients and included any DEG showing significant differences at 0.05 FDR. Mean-centered TMM-normalized counts were used for hierarchical clustering via Ward’s variance minimization method and for comparing the expression of replicative histone genes from 4sU-labeled RNA-seq data. Functional enrichment analyses were carried out using the WebGestaltR package (version 0.4.2). For asynchronous cells, we evaluated the enrichment in Hallmark signatures from MSigDB by over-representation analysis (ORA) for the top 100 upregulated and top 100 downregulated genes, ranked by FDR (Supplementary table 2, sheet 6 and 7). For synchronous cells, we checked enrichment in biological processes associated to specific DEG clusters using Gene Ontology (GO) annotations (Supplementary table 2, sheet 8 and 9). All RNA-seq analyses were carried out with custom Python scripts. pandas (version 0.24.2), NumPy (version 1.16.2) SciPy (version 1.3.1) and fastcluster (version 1.1.26) libraries were used for data manipulation, statistical analysis and hierarchical clustering. matplotlib (version 2.2.4) and seaborn (version 0.9.0) were used for plotting and visualization. R packages were imported into Python using rpy2 (version 2.8.4).

### Analysis of unprocessed replicative histone transcripts from RNA-seq

We used RNA-seq data to investigate the proportion of unprocessed replicative histone transcripts in siCtrl and siASF1 conditions. We retrieved the human SL sequence ‘GGC(C/T)CTTTTCAG(A/G)GCC’ from (Marzluff et al., 2008). Then, we defined a region of 49 nucleotides encompassing the cleavage site, from 20 nucleotides upstream of the SL sequence to 13 nucleotides downstream, reaching the HDE (scheme Figure 4b). Each RNA-seq fragment covering these regions was considered part of an unprocessed transcript. For each replicative histone gene, we compared the total number of fragments mapping to the uncleaved 3’ region to the total number of fragments mapped to the same gene, after removing fragments with discordant paired-end alignments, or a length over 500 nucleotides, or gaps in either of the two mate pairs (i.e. ‘N’ in the CIGAR string). Fragment counts were normalized across samples using DESeq2 median of ratios method (Love et al., 2014). We computed the fraction of unprocessed transcripts by dividing the fragment counts to the uncleaved 3’ region to the total number of fragments mapped to the corresponding replicative histone gene. Then, we compared the relative amounts of unprocessed transcripts in siCtrl and siASF1 from the RNA-seq data in either asynchronous or synchronized cells. A replicative histone gene is considered if its minimal expression across the samples is over 100 reads and the relative amount of unprocessed transcripts is not null in the siCtrl condition.

### Single-molecule RNA FISH

To visualize transcription of the cluster of four replicative histone genes (HIST1H4C, HIST1H1E, HIST1H2BF and HIST1H4E) with single-molecule RNA FISH, a single custom probe set was designed using Stellaris® Probe designer software (LGC, Biosearch Technologies), with cDNA sequences of the replicative histone transcripts used as the input (See probe sequences in supplementary table 1). Single-molecule RNA FISH was performed according to the protocol for adherent cells provided by Biosearch technologies. Briefly, cells were grown on coverslips and following transfections, the cells were fixed with 3.7% formaldehyde in 1X PBS, washed twice with PBS and permeabilised using 70% ethanol. The cells were incubated with 125 nM of each probeset (one for histone genes and second one for a control gene *FOS* in the Stellaris RNA FISH Hybridization Buffer (Biosearch Technologies; SMF-HB1-10) overnight at 37°C in humified chamber. Next day, the coverslips were washed using wash buffers (Biosearch Technologies; SMF-WA1-60, SMF-WB1-20) and 5 ng/ml DAPI was added in the last washing step to counterstain the nucleus. The cells were mounted in the Vectashield Mounting Medium and imaging was done using Zeiss Apotome microscope using 100X objective. Typically, for each condition, 6-7 fields of view (FOVs) were acquired, with each FOV being a stack of 27 z-planes spanning the full nuclear volume with steps of 0.3 μm.

### Analysis of the single-molecule RNA FISH experiments

Nuclei were segmented based on DAPI staining using a custom-written Fiji script and manually curated (elimination of debris, separation of nuclei segmented as one object). Using custom-written Python scripts, RNA spots were detected in the images by detecting local maxima after band-pass filtering, followed by an iterative 3D Gaussian mask fit (Thompson et al., 2002) on the raw images, yielding fluorescence intensities and 3D coordinates of the spots. Fluorescence intensities were then normalized by the intensity of single RNAs, determined as the mode of the intensity distribution observed in segmented nuclei. Sites of transcription were observed as bright spots, each one representing transcription of at least one of the four labeled genes in the cluster. The average intensity of single RNAs was used to estimate the number of RNAs within transcription sites. Consistently with DNA-FISH data from Ghule et al, 2009, we observed at most 3 transcription sites per nucleus. Hence, the total number of RNAs at transcription sites from the labeled genes was approximated as the sum of the 3 brightest spots in each nucleus. Fiji and Python scripts available on https://github.com/CoulonLab/FISHingRod.

### Reverse transcription- quantitative PCR (RT-qPCR)

To obtain the cDNA, reverse transcription PCR was performed using 1 μg of RNA for each sample using the SuperScript^TM^ III First Strand Synthesis SuperMix (Invitrogen, 11752-050) according to the manufacturer’s instructions. We performed quantitative PCR using PowerSYBR Green PCR Master Mix (ABI, 4367659) with three technical replicates in 384-well plates using the QuantStudio 5 Real-Time PCR machine. The relative expression was calculated by ΔΔct method, where the Ct values for PPIA endogenous control are subtracted from each gene of interest (Δct) and then for normalization, we subtracted the Δct of the control (siCtrl) condition for each gene of interest (ΔΔct). Relative expression was calculated as the fold change and plotted using 2^(ΔΔct)^. Primers used in the study are mentioned in supplementary table 1.

### Protein extracts and Western blotting

The cytosolic and nuclear extracts were prepared from HeLa and U2OS cells using the protocol from (Martini et al., 1998). Briefly, cells were lysed with hypotonic buffer using Dounce homogenizer and then centrifuged. The supernatant (cytosolic extract) was collected, and the pellet was treated with lysis buffer containing 300 mM NaCl. After centrifugation, the supernatant (nuclear extract) was collected, and the pellet was treated with benzonase and sonicated. After centrifugation the supernatant containing the chromatin fraction was collected. The extracts were quantified using Pierce BCA protein assay kit (23227). The samples were loaded in various indicated amounts with the 1X loading buffer and DTT on the Bis-Tris 4-12% gel (NuPage) with PageRuler protein ladder. Gels were run at 120 V for ∼1.5 h. Gels were then transferred onto a 0.2 μm nitrocellulose membrane using BioRad Turbo Transblot. Membranes were stained using Pierce reversible protein stain to check the transfer efficiency, blocked in 5% milk prepared in PBST for one hour at room temperature, then incubated in primary antibodies (supplementary table 1) diluted in 5% BSA-PBST at 4°C overnight on a rocker. Membranes were then washed with PBST and incubated in secondary antibodies conjugated with horseradish peroxidase (HRP) with 5% milk PBST for one hour at room temperature. Secondary antibodies were prepared in dilutions of either 1:20,000 (rabbit) or 1:10,000 (mouse, rat). Membranes were then washed with PBST, then exposed with SuperSignal West Dura chemiluminescent reagent. Membranes were imaged using ChemiDoc Touch system and processed with ImageLab software.

## ACKNOWLEDGMENTS

This work was supported by the ERC-2015-ADG-694694 “ChromADICT” and the Ligue Nationale contre le Cancer (Equipe labellisée Ligue). It benefited from ANR-11-LABX- 0044_DEEP and ANR-10-IDEX-0001-02 PSL; ANR-14-CE10-0013 “CELLECTCHIP”; and ANR-16-CE12-0024 “CHIFT”. A.G. was also supported by (H2020 Marie Skłodowska-Curie Actions, grant agreement 798106 “REPLICHROM4D” and ERC-2015-ADG-694694 “ChromADICT”.

## SUPPLEMENTARY FIGURE LEGENDS

**Supplementary figure 1.**
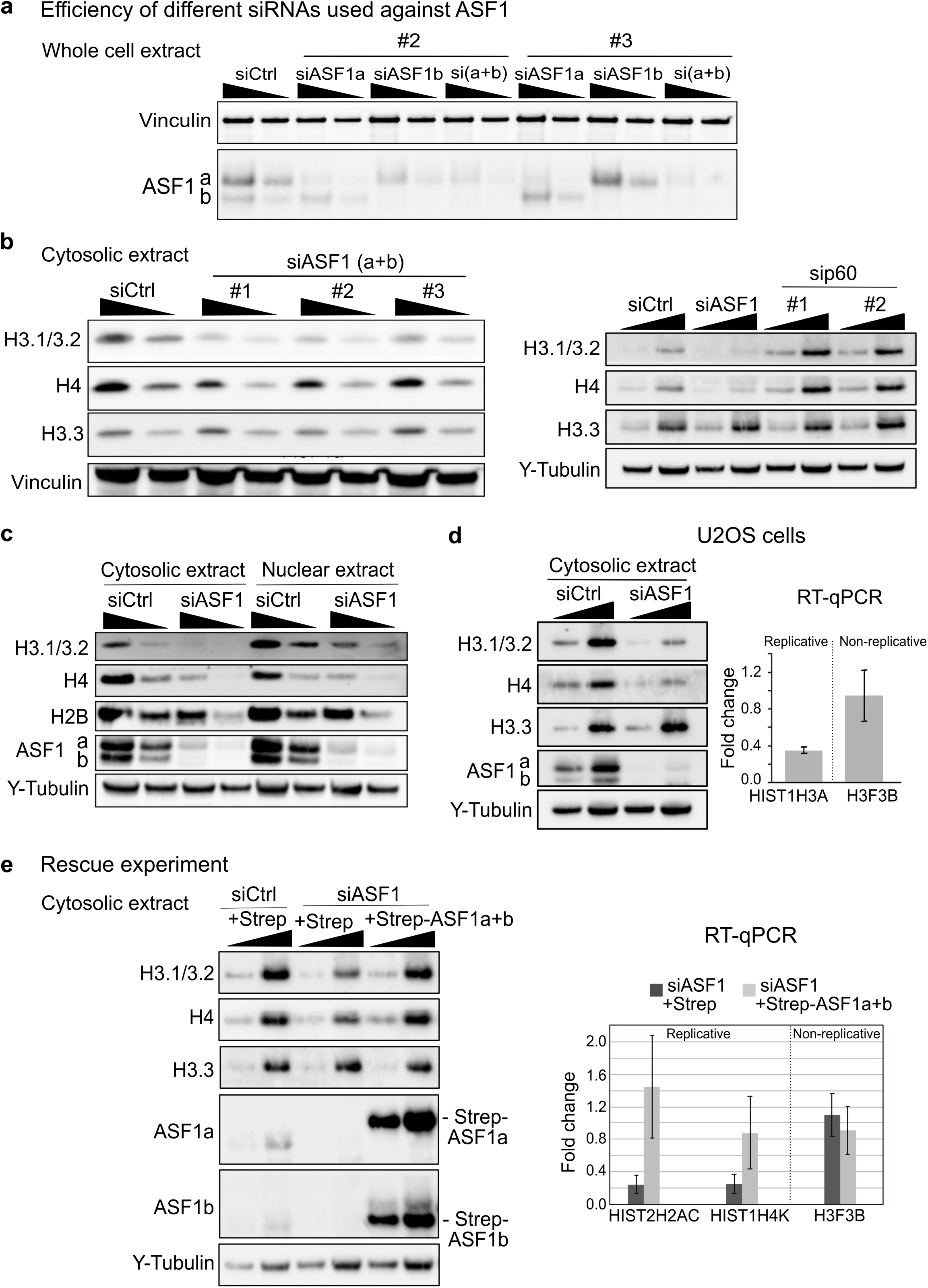
ASF1 depletion reduces replicative histones at the levels of both soluble protein and total RNA. **(a)** Western blot analysis of whole cell extract from HeLa cells harvested 48 h after siRNA transfection (siCtrl, siASF1a (#2 or #3), siASF1b (#2 or #3) or ASF1a+b (#2 or #3)). Two amounts 1x and 2x were loaded. ASF1a and ASF1b were examined, vinculin was used as loading control. **(b) (Left)** Western blot analysis of cytosolic extract from HeLa cells harvested 48 h after siRNA transfection with siCtrl or ASF1a+b (#1, #2 or #3) and **(Right)** with siCtrl, ASF1a+b or sip60 (#1 or #2). Two amounts 1x and 2x were loaded. Replicative histones H3.1/3.2 and H4 and the non-replicative H3.3 were examined, γ-tubulin was used as loading control. **(c)** Western blot analysis of cytosolic and nuclear extracts from HeLa cells harvested 48 h after siRNA transfection (siCtrl or siASF1). Two amounts 1x and 2x were loaded. ASF1a and b, replicative histones H3.1/3.2, H4 and H2B were examined, γ-tubulin was used as loading control. **(d) (Left)** Western blot analysis of cytosolic extract from U2OS cells harvested 48 h after siRNA transfection (siCtrl or siASF1). Two amounts 1x and 2x were loaded. ASF1a and b, replicative histones H3.1/3.2 and H4 and the non-replicative H3.3 were examined, γ-tubulin was used as loading control. **(Right)** RT-qPCR analysis of histone RNA levels in U2OS cells. Total RNA was isolated from U2OS cells harvested 48 h after siRNA transfection (siCtrl or siASF1). Individual histone genes are the replicative HIST1H3A (H3.1) and the non-replicative H3F3B (H3.3). The fold change in transcript levels as compared to siCtrl corresponds to the mean of two independent biological replicate experiments. **(e)** Rescue experiment by overexpression. HeLa cells were transfected with siRNAs (siCtrl or siASF1), then, 24 h later with expression vectors, either with the empty vector pEXPR-IBA105 (Strep) or with those expressing Onestrep-ASF1a and Onestrep-ASF1b (Strep-ASF1a+b). The cells were harvested 48 h after siRNA transfection and either proteins or total RNA were prepared. **(Left)** Western blot analysis of cytosolic extract. Two amounts 1x and 2x were loaded. ASF1a and ASF1b, replicative histones H3.1/3.2 and H4 and the non-replicative H3.3 were examined, γ-tubulin was used as loading control. **(Right)** RT-qPCR analysis. Individual histone genes are the replicative HIST2H2AC (H2A type 2-C) and HIST1H4K (H4) and the non-replicative H3F3B (H3.3). The fold change in transcript levels as compared to siCtrl+Strep corresponds to the mean of two independent biological replicate experiments.

**Supplementary figure 2.**
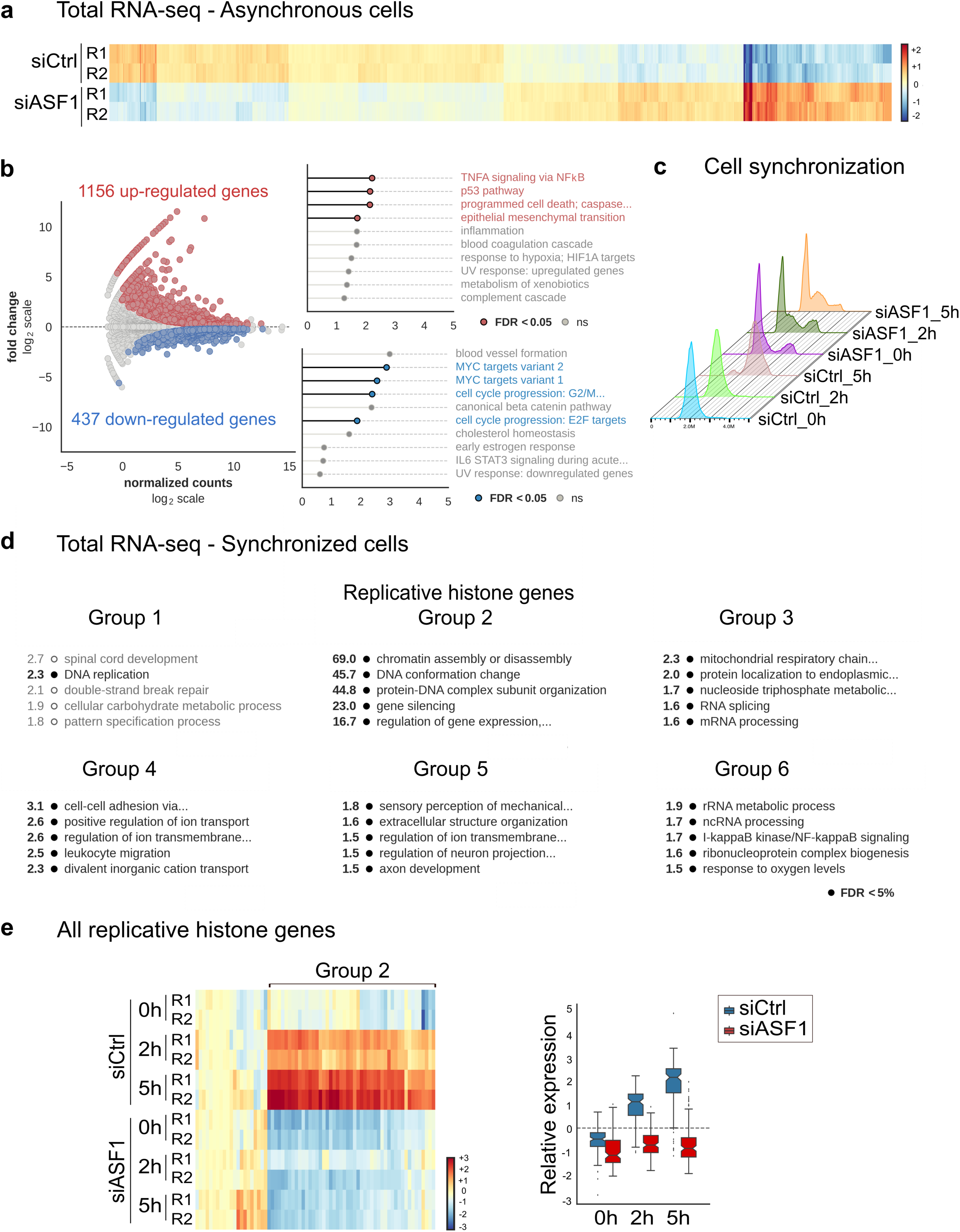
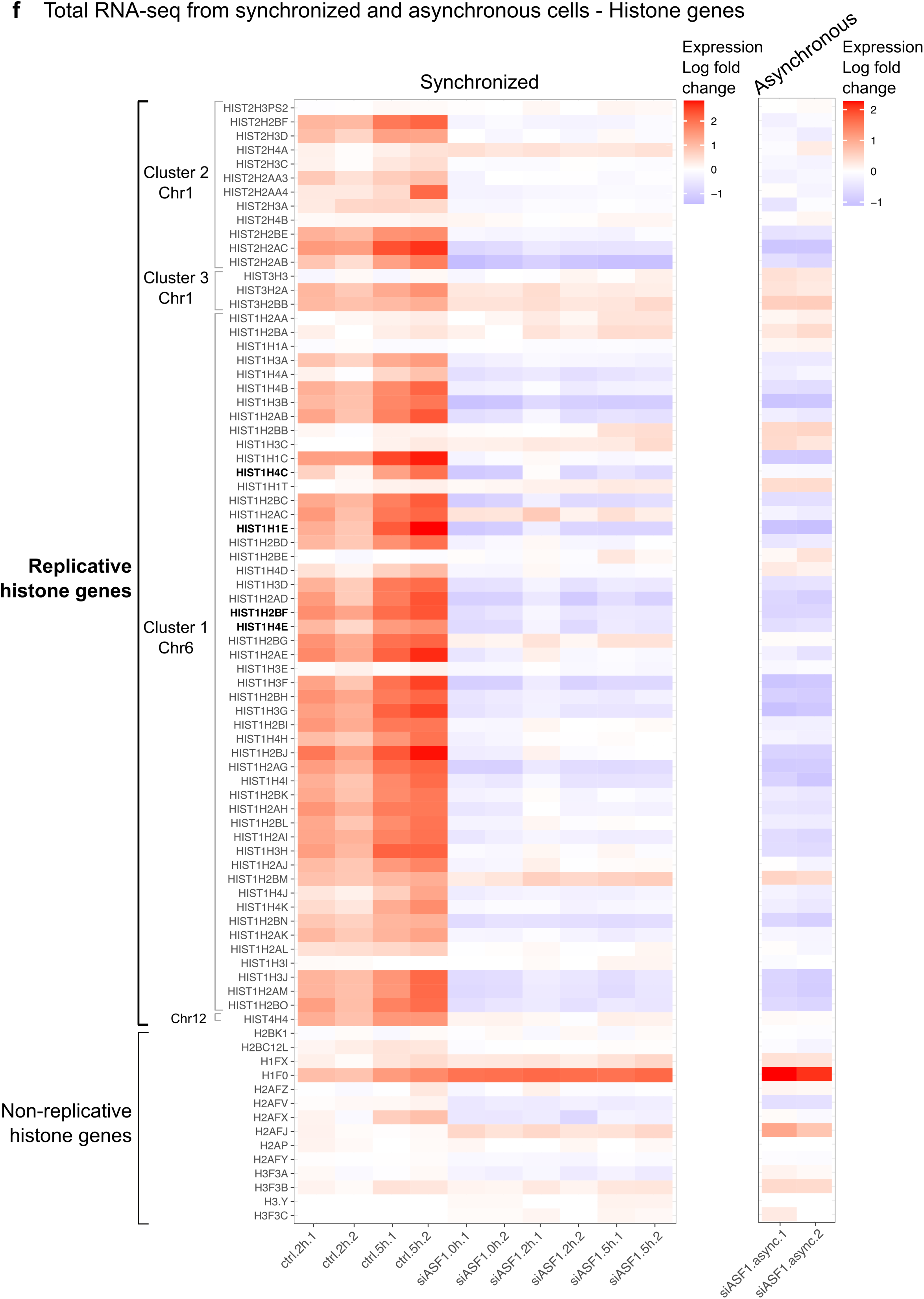
Impact of ASF1 depletion on transcriptome - RNA-seq data from asynchronous and synchronized HeLa cells in siCtrl and siASF1 conditions. **(a)** Heat map showing hierarchical clustering of samples (rows) and differentially expressed genes (columns) between siCtrl and siASF1 conditions in asynchronous cells, including all genes showing significant differences at FDR < 0.05. The color gradient is proportional to the expression of each gene in a given sample, relative to their average across all samples (log_2_ normalized counts, mean-centered): from blue (lower than average) to red (higher than average). The experiment was performed in duplicate, replicate 1 (R1) and replicate 2 (R2). **(b) (Left)** MA plot showing log_2_ fold-change between siASF1 and siCtrl versus baseline expression levels across all genes. Differentially expressed genes at FDR < 0.05 are highlighted in blue (down-regulated) or red (up-regulated). **(Right)** Hallmark signatures associated with the top 100 up-regulated (top panel) or down-regulated genes (bottom panel), as identified by over-representation analysis (ORA) with WebGestaltR v0.4.2. Signatures showing significant enrichment at FDR < 0.05 are highlighted in red (for up- regulated genes) and blue (for down-regulated genes). **(c)** Cell synchronization. FACS analysis of the different conditions. **(d)** Six groups of genes identified by hierarchical clustering from differential gene expression between siASF1 and siCtrl in synchronized HeLa cells (see figure 2b and 2d). For all groups of genes, the top 5 Gene Ontology (GO) annotations are shown (over- representation analysis with WebGestaltR, v0.4.2). GO terms showing significant enrichment at FDR < 0.05 are highlighted in black. Group 2 corresponds only to replicative histone genes. **(e) (Left)** Heatmap showing hierarchical clustering of all replicative histone genes in siCtrl and siASF1 conditions. Color gradient shows expression relative to average as in 2b (log_2_ normalized counts, mean-centered): from blue (lower than average) to red (higher than average). **(Right)** The box plots show the distribution of relative expression levels of all replicative histone genes in siCtrl (in blue) and siASF1 (in red) conditions at 0 h (G1/S), 2 h (early S) and 5 h after release from the G1/S block (log_2_ normalized counts, mean-centered across all samples and averaged by experimental condition). **(f)** Heat maps showing histone gene expression from the RNA-seq data in the two replicates. Log fold change of expression for each of the expressed histone genes in the siCtrl and siASF1 conditions from RNA-seq data in asynchronous and synchronized HeLa cells. The log fold change was computed in the samples of asynchronous cells in siASF1 condition according to the samples of asynchronous cells in siCtrl condition in respect of the replicate. For the samples of synchronized cells in S-phase, their log fold changes were computed according to the sample of siCtrl condition with cells arrested at G1/S checkpoint (’’0h’’) of the same replicate. The replicative histone genes are displayed first according to their genomic order in each cluster. In bold are shown the genes used as probes for single-molecule RNA FISH. Non-replicative histone genes are at the bottom of the panel ordered by their type of histone.

**Supplementary figure 3.**
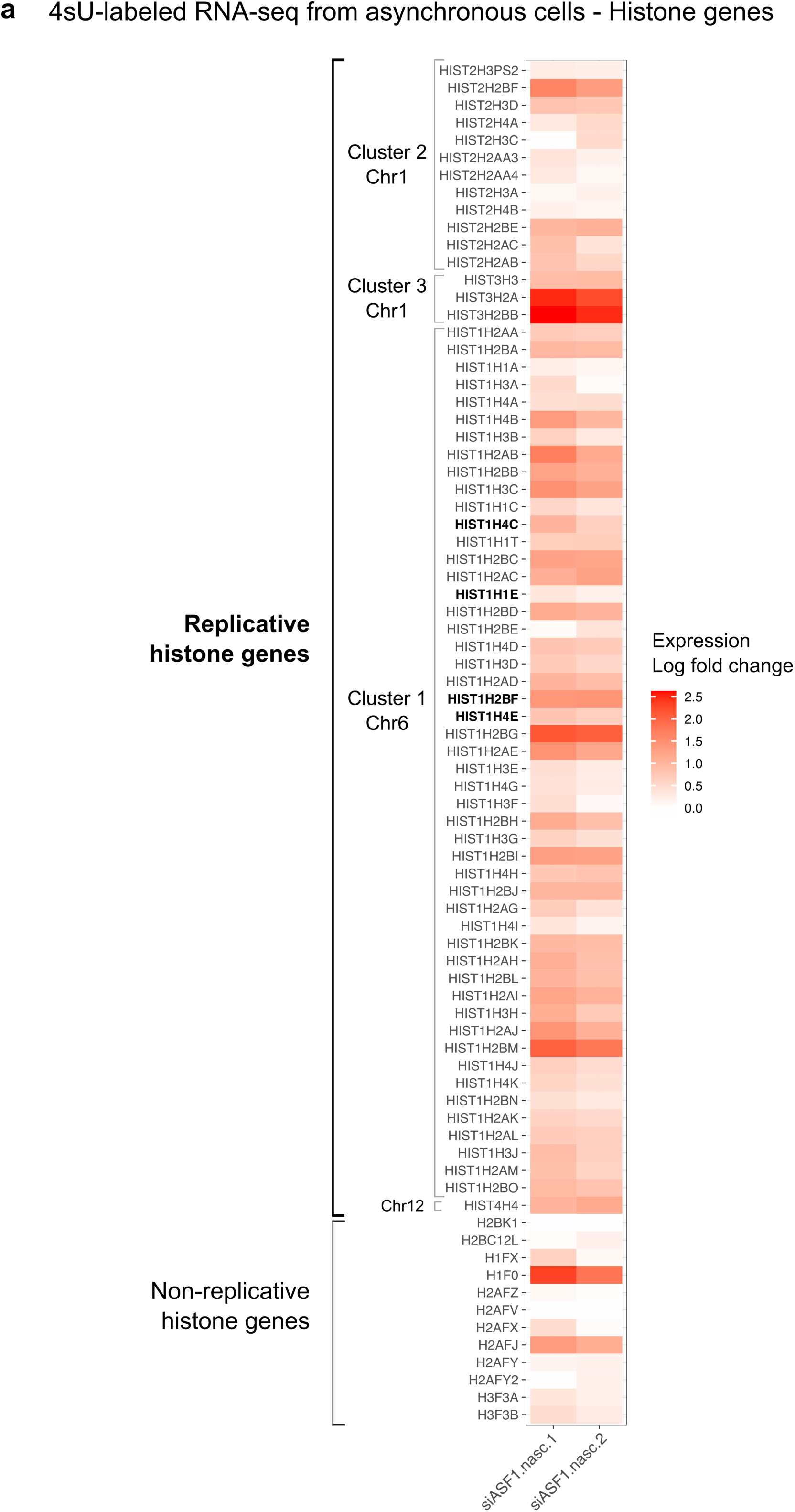
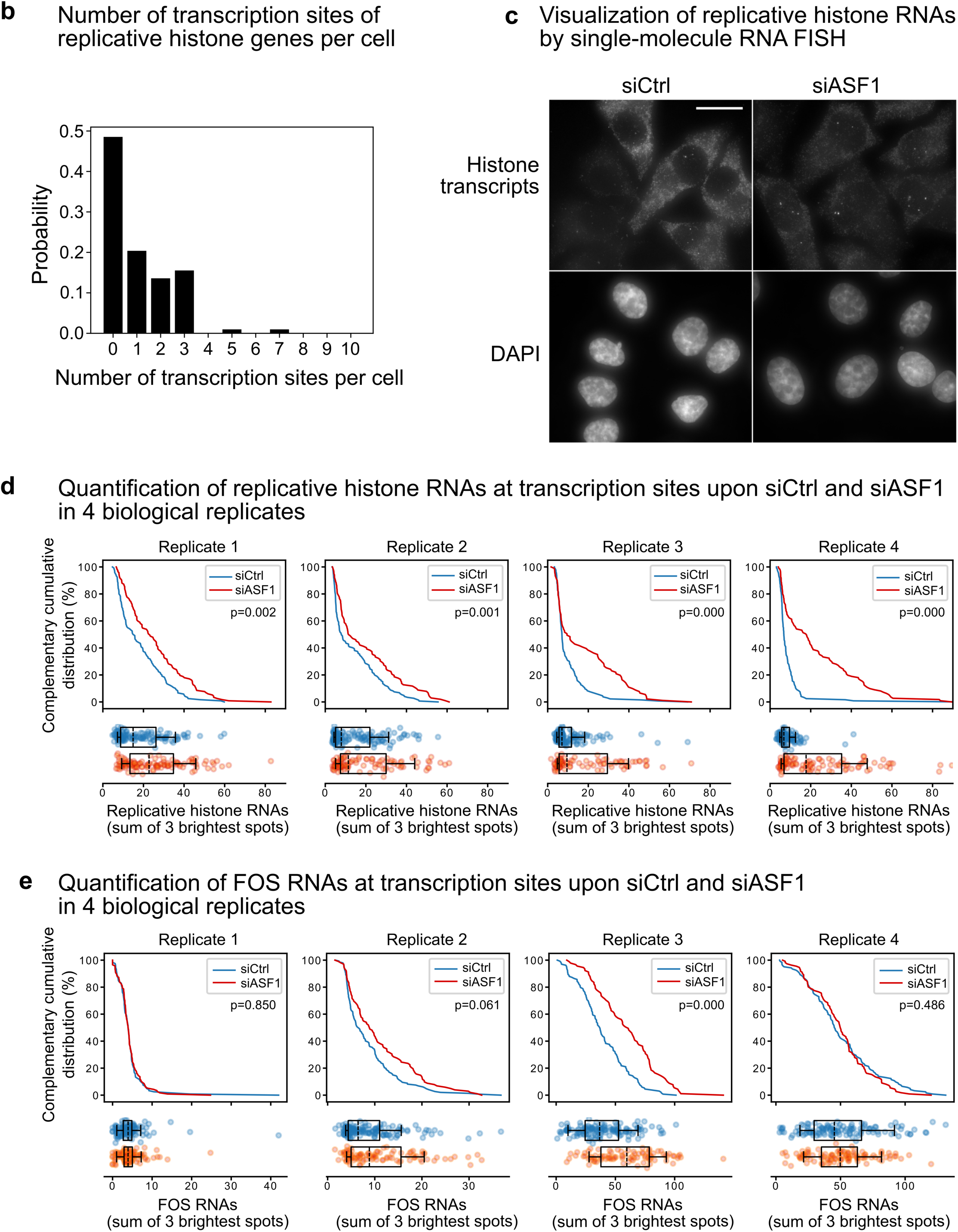
ASF1 depletion increases replicative histone nascent RNAs at transcription sites. **(a)** Heat map showing histone gene expression from the 4sU-labeled RNA sequencing data in the two replicates. The log fold change was computed in the samples of cells in siASF1 condition according to the samples of cells in siCtrl condition in respect of the replicate. The replicative histone genes are displayed first according to their genomic order in each cluster. In bold are shown the genes used as probes for single-molecule RNA FISH. Non-replicative histone genes are at the bottom of the panel ordered by their type of histone. **(b)** Histogram showing the number of transcription sites of replicative histone genes per cell. **(c)** Representative images of single-molecule RNA FISH for replicative histones in siCtrl and siASF1 conditions. **(d)** Complementary cumulative distributions of the number of replicative histone RNAs at transcription sites per nucleus (calculated as the sum of the 3 brightest spots) in control condition (siCtrl, blue) and upon ASF1 depletion (siASF1, red). Scatterplots show the numbers of replicative histone RNAs at transcription sites per nucleus in control condition (siCtrl, blue) and upon ASF1 depletion (siASF1, red). Dashed lines show the median RNA numbers, boxes show the top and bottom quartiles and whiskers show the 10- and 90-percentile. The p-values of Kolmogorov-Smirnov test are indicated for each replicate. All four individual biological replicates are shown. **(e)** Complementary cumulative distributions of the number of FOS RNAs at transcription sites per nucleus (calculated as the sum of the 3 brightest spots) in control condition (siCtrl, blue) and upon ASF1 depletion (siASF1, red). Scatterplots show the numbers of FOS RNAs at transcription sites per nucleus in control condition (siCtrl, blue) and upon ASF1 depletion (siASF1, red). Dashed lines show the median RNA numbers, boxes show the top and bottom quartiles and whiskers show the 10- and 90-percentile. The p-values of Kolmogorov-Smirnov test are indicated for each replicate. All four individual biological replicates are shown.

**Supplementary figure 4.**
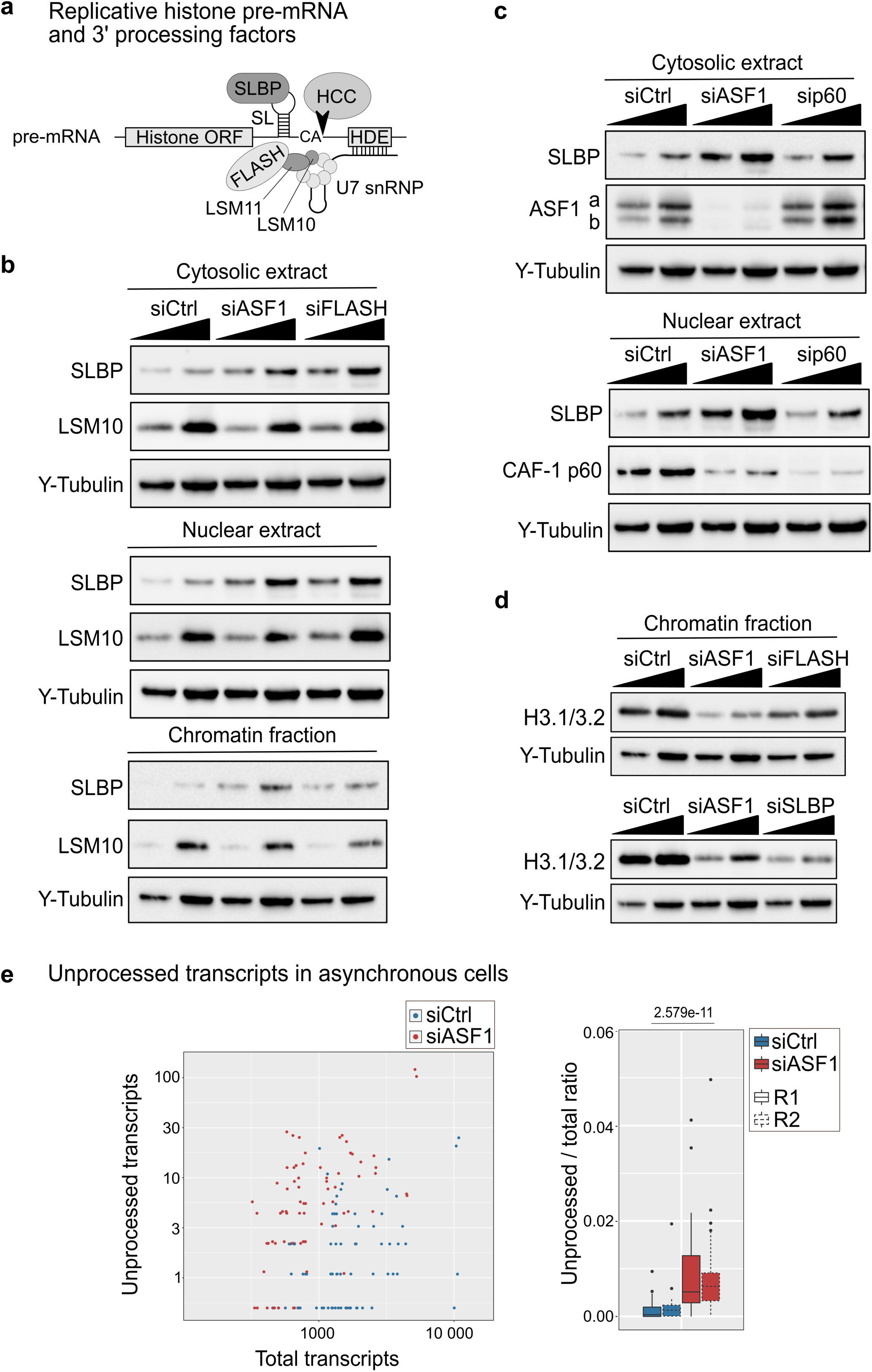
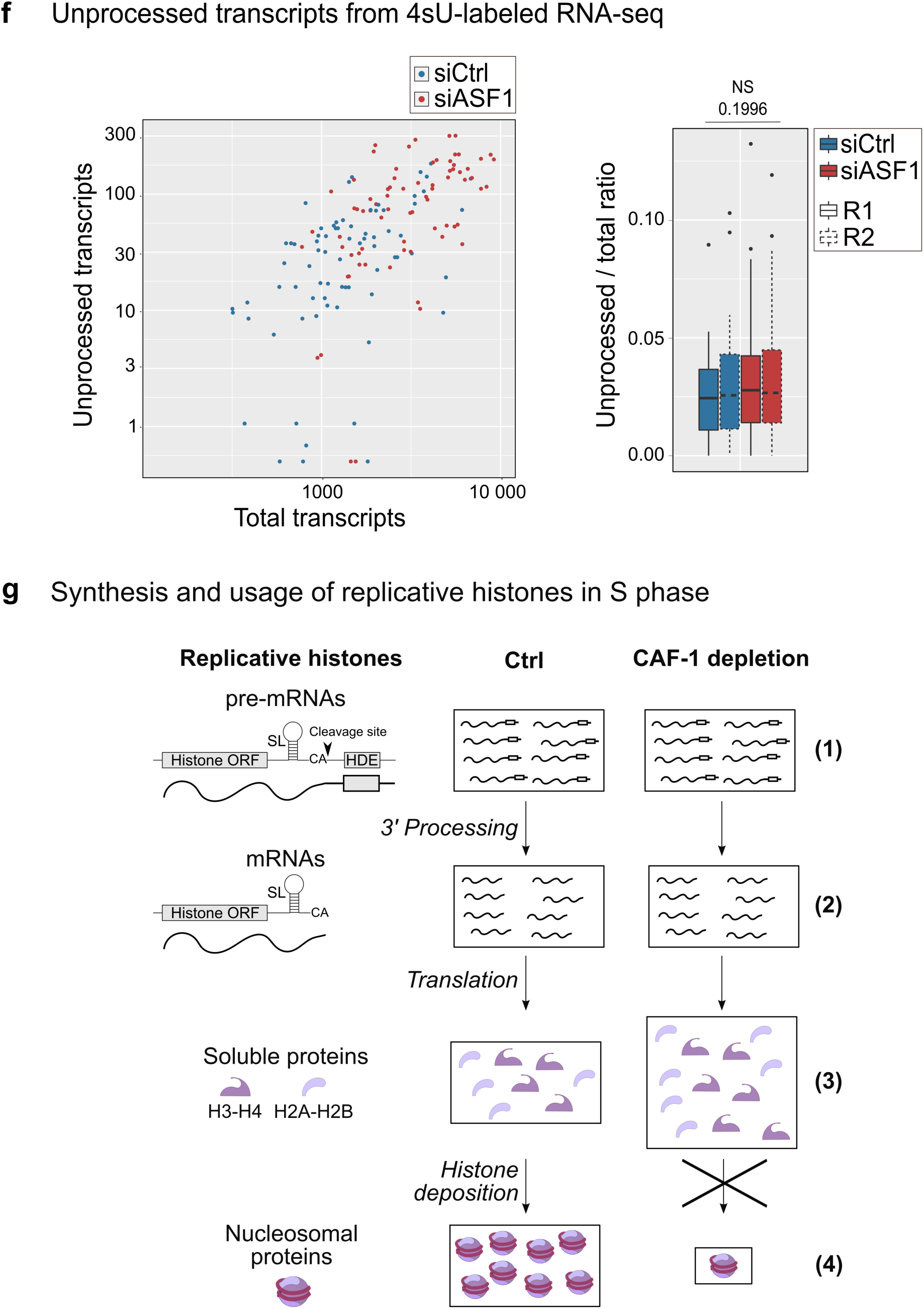
ASF1 depletion affects the 3’ processing of replicative histone transcripts. **(a)** Scheme showing the structure of a replicative histone pre-mRNA and the main known 3’ processing factors (Marzluff and Koreski, 2017; Mendiratta et al., 2019). These 3’ processing factors are Stem loop binding protein (SLBP), FLice-associated huge protein (FLASH), U7 small nuclear ribonucleoprotein (snRNP) and the Histone cleavage complex (HCC). U7-snRNP is composed of a U7 snRNA that binds the HDE and of a heptameric ring made of five Sm proteins and two Sm-like proteins LSM10 and LSM11. **(b)** Western blot analysis of cytosolic, nuclear and chromatin fractions from HeLa cells harvested 48 h after siRNA transfection (siCtrl, siASF1 or siFLASH). Two amounts 1x and 2x were loaded. SLBP and LSM10 were examined, γ-tubulin was used as loading control. **(c)** Western blot analysis of cytosolic and nuclear extracts from HeLa cells harvested 48 h after siRNA transfection (siCtrl, siASF1 or sip60). Two amounts 1x and 2x were loaded. SLBP was examined, γ-tubulin was used as loading control. **(d)** Western blot analysis of chromatin fraction from HeLa cells harvested 48 h after siRNA transfection (siCtrl, siASF1, siFLASH or siSLBP). Two amounts 1x and 2x were loaded. The replicative H3.1/3.2 was examined and γ-tubulin was used as loading control. **(e)** Comparison of the amounts of unprocessed transcripts between siASF1 and siCtrl conditions using our RNA-seq data of asynchronous cells. **(Left)** The replicative histone genes are plotted according to their normalized number of unprocessed and total transcripts. Every dot is one replicative histone gene. Genes with a null number of unprocessed transcripts are displayed at the value of 0.5. **(Right)** The box plots show the ratio of the number of unprocessed transcripts to total transcripts (total number of fragments mapping in each annotated replicative histone gene). The results for the two replicates R1 and R2 are shown. The p-value is indicated. **(f)** Comparison of the amounts of unprocessed transcripts between siASF1 and siCtrl conditions using our 4sU-labeled RNA-seq data. **(Left)** The replicative histone genes are plotted according to their normalized number of unprocessed and total transcripts. Every dot is one replicative histone gene. Genes with a null number of unprocessed transcripts are displayed at the value of 0.5. **(Right)** The box plots show the ratio of the number of unprocessed transcripts to total transcripts (total number of reads mapping in each annotated replicative histone gene). The results for the two replicates R1 and R2 are shown. The p-value is indicated. **(g)** Model showing the impact of p60 CAF-1 depletion on the synthesis and usage of replicative histones during S phase. In contrast to ASF1 depletion (see model figure 4d), CAF-1 depletion does not affect the replicative histone RNA metabolism (1,2) but leads to an accumulation of soluble replicative histones (3) most likely due to a defect of usage (histone deposition) reflected by a decrease of nucleosomal replicative histones (4). We propose that upon CAF-1 depletion the excess of soluble histones could be buffered by ASF1 in a manner that could compare to the buffering that we reported for the excess of soluble replicative histones upon replicational stress (Groth et al., 2005).

## Notes

### Competing Interest Statement

The authors have declared no competing interest.

