## Supplementary Table 1 for "Regulation of replicative histone RNA metabolism by the histone chaperone ASF1"

| **Antibodies** | **Company** | **Catalog number or reference** |
|---|---|---|
| anti-ASF1a | Agrobio | Groth et al., 2005 |
| anti-ASF1a | Cell Signaling | #2990 |
| anti- ASF1b | Agrobio | Groth et al., 2005 |
| anti- H3.1/2 | Active Motif | 61629 |
| anti-H3.3 | Millipore | 09-838 |
| anti-H4 | Abcam | ab31830 |
| anti-H2B | Abcam | ab1790 |
| anti-p60 (CAF-1) | Agrobio | Groth et al., 2005 |
| anti-SLBP | Abcam | ab181972 |
| anti-LSM10 | Abcam | ab180128 |
| anti-γ-Tubulin | SIGMA | T5326 |
| anti-Vinculin | SIGMA | V9131 |

| **siRNA** | **Sequence** | **Reference** |
|---|---|---|
| siCtrl | UGGUUUACAUGUCGACUAA | |
| siAsf1a#1 | GUGAAGAAUACGAUCAAGUUU | |
| siASF1b#1 | CAACGAGUACCUCAACCCUUU | |
| siAsf1a#2 | GAGACAGAAUUAAGGGAAAUU | |
| siAsf1b#2 | CGGACGACCUGGAGUGGAAUU | |
| siAsf1a#3 3'UTR | CCUGAAAUUCCGUAAGUAUU | |
| siAsf1b#3 3'UTR | CCUUGAGUACCAUUGAUCUU | |
| siFLASH | CCGCAAGGAUGAAGAAAUAUU | Mejlvang et al., 2013 |
| siSLBP#1 | GGAUGUGAUUUGCAAGAAAUU | Mejlvang et al., 2013 |
| sip60#1 | GCGUGUGGCUUUCAAUGUUUU | Polo et al., 2006 |
| sip60#2 | UCUUGCUCGUCAUACCAAAUU | Polo et al., 2006 |

**RNA FISH probes**

| Name | Sequence | Name | Sequence |
|---|---|---|---|
| FOS_1 | cagccactgcttttataaca | H4C5_1 | ccgggaattcggacctaaat |
| FOS_2 | agatgcggttggagtacgag | H4C5_2 | gcgacggtagtattcttata |
| FOS_3 | cagtcttggcttctcagatg | H4C5_3 | catgaccaacacaccaacga |
| FOS_4 | aacatcatcgtggcggttag | H4C5_4 | tggatgttatctcgcaggac |
| FOS_5 | ctcgtagtctgcgttgaagc | H4C5_5 | ggatggcaggcttggtaatg |
| FOS_6 | gagtggtagtaagagaggct | H4C5_6 | cagaaatgcgcttgacaccc |
| FOS_7 | catgctggagaaggagtctg | H4C5_8 | gtcacagcatcacgaatcac |
| FOS_8 | ggaatgaagttggcactgga | H4C5_9 | ctgtcactgtcttgcgtttg |
| FOS_9 | ctggtcgagatggcagtgac | H4C5_10 | tgtctcttcagcgcgtagac |
| FOS_10 | cgaaagggtgaggggctctg | H1.4_1 | gtgagcgagagcaattcgag |
| FOS_11 | ctgtcatggtcttcacaacg | H1.4_2 | ctcggacatgttgaaggcaa |
| FOS_12 | ttctcatcttctagttggtc | H1.4_3 | ttcttcttcacgggagtctt |
| FOS_13 | aatctcggtctgcaaagcag | H1.4_4 | gctttagtaatgagctcgga |
| FOS_14 | gtcatcagggatcttgcagg | H1.4_5 | ttgagagcggccaaagatac |
| FOS_15 | cagacatctcttctgggaag | H1.4_6 | tcttgagacccagcttgatg |
| FOS_16 | ccagtcagatcaagggaagc | H1.4_7 | tgagtttgaaggaacccgac |
| FOS_17 | ctcagggtcattgaggagag | H1.4_8 | ctttttagccttaggcttgg |
| FOS_18 | ctgatgctcttgacaggttc | H1.4_9 | cttctttggggtcttcttgg |
| FOS_19 | caggaagtcatcaaagggct | H1.4_10 | tcgggcttttcgcttttttg |
| FOS_20 | agataggtccatgtctggca | H1.4_11 | ttttttggcttggctgcttt |
| FOS_21 | agtctgctgcatagaaggac | H1.4_12 | ctttctactttttcttggct |
| FOS_22 | ctgagcgagtcagaggaagg | H1.4_13 | cttctaagcagttggccaaa |
| FOS_23 | tacaggaaccctctagggaa | H2BC7_1 | cacttcgtccacaacctaaa |
| FOS_24 | gtttcacgcacagataaggt | H2BC7_2 | gcaggttcaggcatgataaa |
| FOS_25 | gatgctttcaagtccttgag | H2BC7_3 | ttttggagcaggagcggact |
| FOS_26 | gtaaggacttgagtccacac | H2BC7_4 | gggattgggtatgaagacgt |
| FOS_27 | ttttgctacatctccggaag | H4C3_1 | gaccagacatgattcctatc |
| FOS_28 | cagctctctgaagtgtcact | H4C3_2 | cttacgatggcgcttagcac |
| FOS_29 | ggctcaacatgctactaact | H4C3_3 | tgtaatgccctggatgttat |
| FOS_30 | cccaatagattagttaatgc | H4C3_4 | Cgagccaaacggcgaatagc |
| FOS_31 | gcaccaggttaattccaata | H4C3_5 | Tcctcatagataagaccgga |
| FOS_32 | cactattgccaggaacacag | H4C3_6 | Aaaccttaagcacacctcga |
| FOS_33 | gtcgtatcttttcttagtat | H4C3_7 | Tgacggcgtctcgaataacg |
| FOS_34 | ctcaacaatgcatgatcagt | H4C3_8 | Ctgtgacagttttgcgcttg |
| FOS_35 | aatgtcagaacattcagacc | H4C3_9 | Agggcatatactacatccat |
| FOS_36 | tccacatgtcaaaagacctc | H4C3_10 | Cgaagccatacagagtgcgc |
| FOS_37 | gtcgcattcaacttaaatgc | H4C3_11 | Agaccgcgtattcttagatt |

**RT-qPCR primer sequences**

| Genes | Forward primer | Reverse primer |
|---|---|---|
| HIST2H2AC | GCTCGGGACAACAAGAAGAC | ATTTGCTTTTGGCTTTGTGG |
| HIST1H4K | TACTGCGCGACAATATCCAG | CAACCACCGAAACCGTAGAG |
| HIST1H1E | TTCAACATGTCCGAGACTGC | AGGCGGCAACAGCTTTAGTA |
| HIST1H3A | AGATCCGCCGTTATCAGAAGTC | CAGGCGCTGGAAAGGTAGTT |
| CENPA | TGGACTTCAATTGGCAAGCC | AGTAACTCGGCCTGCATGTA |
| H3F3A | TAAAGCACCCAGGAAGCAAC | AGGGAAGTTTGCGAATCAGA |
| H3F3B | AAGCTGCCCTTCCAGAGGTT | ACCTCAGGTCGGTTTTGAAATC |
| PPIA | CATCTGCACTGCCAAGACTGA | TTCATGCCTTCTTTCACTTTGC |
